# Exercise Evokes Retained Motor Performance without Neuroprotection in a Mouse Model of Parkinson’s Disease

**DOI:** 10.1101/2024.09.20.614034

**Authors:** Henry M. Skelton, Nathaniel Hyman, Alejandra M. Fernandez, Emma Acerbo, Madison Scott, Ken Berglund, Claire-Anne Gutekunst, Robert E. Gross

## Abstract

Exercise has been extensively studied in Parkinson’s Disease, with a particular focus on the potential for neuroprotection that has been demonstrated in animal models. While this preclinical work has provided insight into the underlying molecular mechanisms, it has not addressed the neurophysiological changes during exercise. Here, first, we tested for neuroprotective effects of adaptive wheel running exercise in the 6-hydroxydopamine mouse model of Parkinson’s disease, assessing for dopaminergic cell preservation. Finding none, despite running performance that equaled the pre-parkinsonian state, we probed the neurophysiology of running exercise as a transient state of high motor function amidst an unameliorated Parkinsonian lesion. Exercise was associated with characteristic, excitatory changes in the dopamine-depleted substantia nigra, which could be suppressed along with running itself by dopamine receptor blockade. Going forward, the functional state evoked by exercise merits further study, as it has parallels in human disease and may represent an optimal physiologic target for neuromodulation.

## Introduction

Exercise is associated with a reduced risk of developing Parkinson’s Disease (Sasco et al., 1992; Xu et al., 2010; Yang et al., 2015) as well as slower symptomatic progression (Rafferty et al., 2017; Tsukita et al., 2022), with some thought that it may provide for otherwise elusive disease course modification. In support of that, neuroprotection has been directly demonstrated in rodent models of PD, with the compelling, albeit not universal, finding that exercise reduces dopaminergic cell loss (Ferreira et al., 2021; Hou et al., 2017; Palasz et al., 2019).

A wide range of biomolecular pathways have been implicated in the therapeutic effects of exercise. Among those, the involvement of brain-derived neurotrophic factor (BDNF) signaling (Gerecke et al., 2012; Real et al., 2013; Tajiri et al., 2010; Wu et al., 2011) stands out. As BDNF release is characteristically driven by neuronal activity (Balkowiec and Katz, 2002), the neurophysiology of exercise could play a key role in mediating its impact on PD. If that role is causal, it may even be possible to engage the same mechanisms with neuromodulation, which is an established symptomatic treatment for PD in the form of deep brain stimulation (DBS). Interestingly, there is evidence in rodents that DBS may have clinically unrealized, BDNF-dependent neuroprotective properties (Fischer et al., 2017; Fischer and Sortwell, 2019).

However, the active neuronal changes during exercise have not been well studied in general, nor at all in the context of preclinical PD models. This likely reflects the difficulty of performing neurophysiologic assays during exercise. Neither spontaneous exercise, which occurs sporadically throughout the night in the home cage, nor forced treadmill exercise, which commonly relies on aversive electrical stimulus, are well suited for sensitive physiological measurements that require tethered attachment to complex recording systems.

The challenges of studying exercise in PD models are compounded by the motor deficits inherent to those models. The 6-hydroxydopamine (6-OHDA) mouse model is arguably the most well demonstrated tool for dissecting the mechanisms of circuit therapies (Gradinaru et al., 2009; Schor et al., 2022; Spix et al., 2021), providing for consistent, behaviorally-evident dopamine depletion in a species with many transgenic lines. However, relatively few exercise studies use 6-OHDA mice (Ferreira et al., 2021), possibly because the motor impairment that makes it a valid model system also tends to limit exercise performance.

Previously, we developed adaptive wheel exercise (Skelton et al., 2025), which effectively invokes running in mice, even in the early aftermath of a 6-OHDA lesion. Here, using that adaptive wheel platform and disease model, we tested whether and along what time course exercise would protect against nigral dopaminergic cell loss. Finding no evidence of exercise-based neuroprotection, we were surprised to find that the intractable lesion did not limit exercise itself, and we probed the dopamine-dependence and electrophysiology of that retained motor performance. Wheel running remained dopamine dependent even after unilateral 6-OHDA lesions and was associated with a transient elevation of SNr firing rates.

## Results

### Adaptive Wheel Exercise in 6-OHDA Lesioned Mice

In order to identify the critical period for neuroprotection with exercise in 6-OHDA-lesioned mice, we designed an exercise intervention variably applied across the disease course. We used adaptive wheel exercise to induce running before and as soon as one day after unilateral 6-

OHDA lesioning. As previously described (Skelton et al., 2025), this uses positional tracking to accelerate mice to the limit of their running capacity, rapidly slowing down if the mouse falls off pace to permit recovery (Fig 1A). On each day of exercise intervention, mice were either treated with 40 minutes of adaptive exercise or placed in a locked wheel for the same period of time. Subjects were divided into four groups that were, respectively, exercised only before the lesion (Exercise-Before group, n=7), only after (Exercise-After group, n=7), both before and after (Exercise-Both group, n=8), and a final control group (Sedentary group, n=7) that was sedentary at all time points (Fig 1B). These sessions (with typical exercise illustrated in figure 1C) started four weeks before and ended four weeks after 6-OHDA lesioning (Fig 1D). Due to 6-OHDA related mortality, 28% of all subjects died between days 7 and 14 (Fig 1E), with no significant difference between groups (Fisher’s exact test). Five mice in all of the exercised groups, and six in the sedentary group, reached the endpoint for tissue analysis at 28 days. All analyses are confined to those mice, except as otherwise stated.

**Figure 1.**
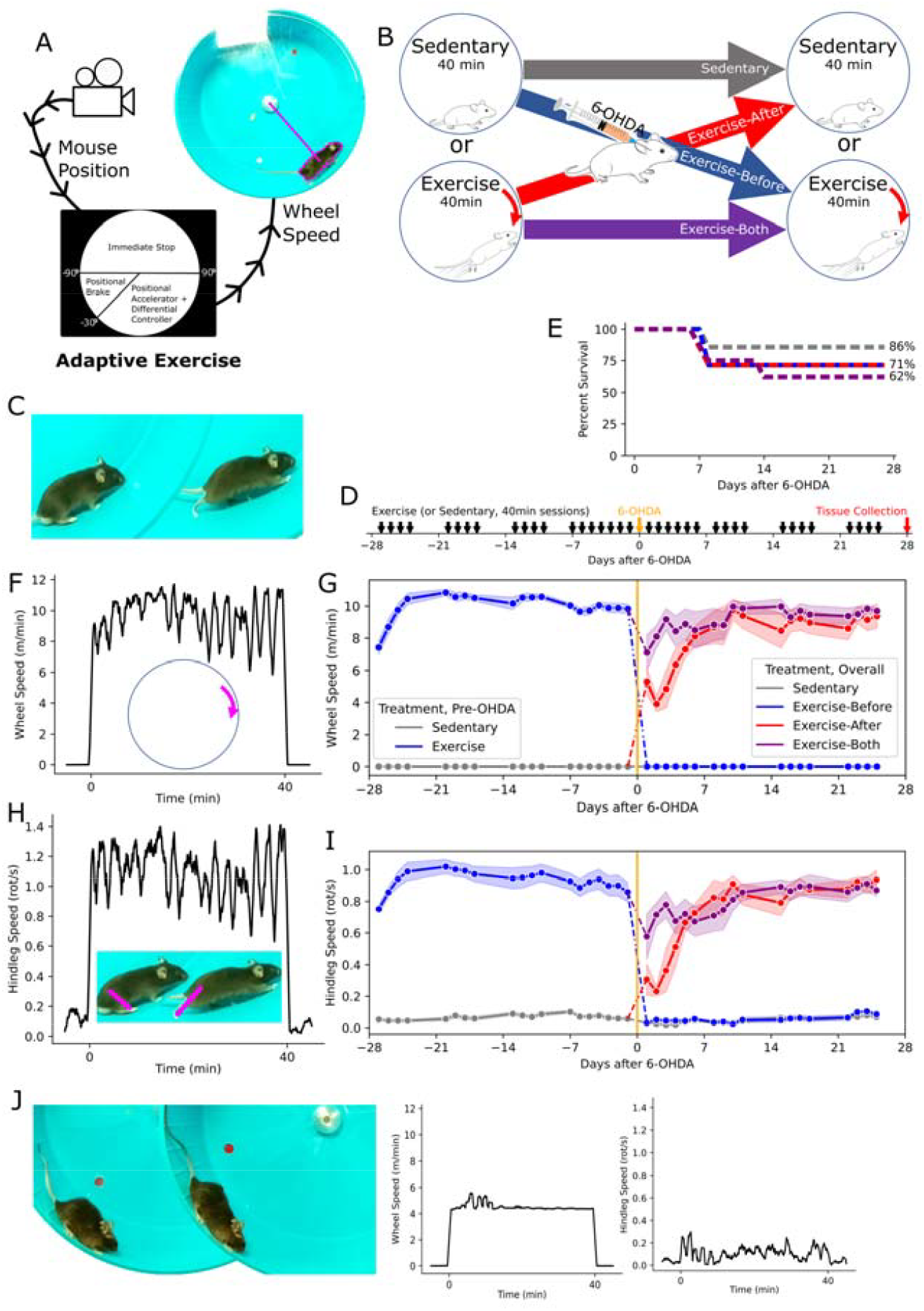
Exercise in 6-OHDA Lesioned Mice. **(A)** Schematic of closed-loop control in exercise, showing tracking based on color thresholding. (**B)** Experimental design, with groups divided according to whether they were exercised or sedentary before and after the 6-OHDA lesion. **(C)** Example of typical running. **(D)** Timeline of the intervention around the 6-OHDA lesion. **(E)** The percentage of surviving mice in each group over the post-lesion time course, with the final percentages indicated. **(F)** Representative tracing of wheel speed, over the course of an adaptive exercise experiment. **(G)** Daily wheel speed for group across the course of the intervention, showing the mean (dots) and SEM (bands). Here, the mice are divided into two groups before the 6-OHDA lesion (day 0), and subdivided into four groups thereafter. **(H)** Representative tracing of hindleg speed, over the course of an adaptive exercise experiment. **(I)** Mean daily hindleg speed, shown as in G. **(J)** Representative image of failure to run, most prevalent in the early post-OHDA period, accompanied by traces of wheel and hindleg speed.

In practice, the adaptive wheel program produced a pattern of high-speed intervals (Fig 1F) with a pre-lesional, session-wise mean wheel speed of 10.0 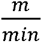 (SD 1.2, Fig 1G). In order to quantify behavioral engagement with this intervention, we measured hindleg motion via markerless pose tracking of video recordings (Fig 1H). Prior to the 6-OHDA lesion, the mean hindleg speed was 0.93 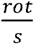 (SD 0.18, Fig 1I). As the locked wheel did not preclude movement, hindleg speed was nonzero, but minimal in the fixed-wheel sessions, with a session-wise mean of 0.07 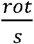 (SD 0.04).

Among mice exercised after the lesion, running performance was impaired in the early period, with a tendency to slide (Fig 1J) rather than run. This was more prevalent in the animals newly introduced to exercise After the lesion, but eventually resolved in all cases.

### Exercise and Hemiparkinsonian Behavior

We measured behavior throughout the experiment, using open-field behavioral recordings quantified via pose tracking (Fig 2A). Once per week, on non-interventional days, we performed recordings immediately after administration of methamphetamine to aggravate hemiparkinsonism (Fig 2B). In the pre-lesion period (through day −14), there was no difference between mice that were exercised and those that were sedentary, nor any change over time, or interaction effect (Mixed ANOVA). In the peri-lesion period (day −14 through 28), there remained no difference between treatment groups or interaction effect, however, there was a significant change over time as hemiparkinsonian ipsiversive turning developed in all groups (p=1.6×10^−14^, Mixed ANOVA). We compared all subjects across days, to assess both the effect of the lesion and progression over time, finding amphetamine induced rotations to be significantly higher on all post-lesion days compared to the last pre-lesion measurement on day −14 (p<0.001, paired t-tests, with Bonferroni correction), with no significant differences among the post-lesion measurements.

**Figure 2.**
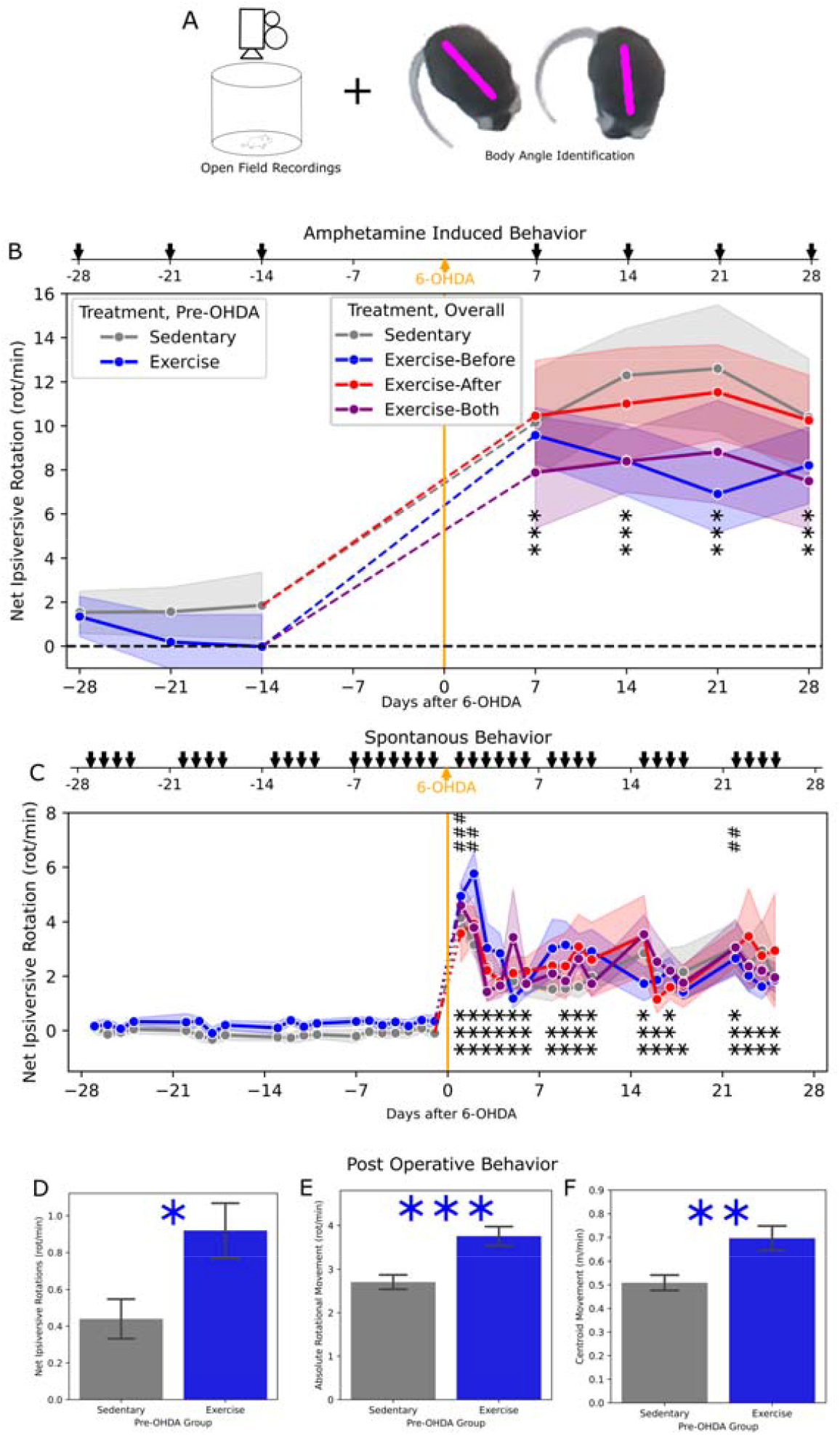
Hemiparkinsonian Behavior. **(A)** Schematic of behavioral testing using open field recordings. **(B)** Measurements of net ipsiversive (relative to the 6-OHDA injection) rotation performed immediately after injection of methamphetamine, shown as the groupwise mean (dots) and SEM (bands). With no group differences, significance is indicated for the aggregate difference between each day and the last pre-OHDA measurement (day-4) by pairwise t-test. **p< 0.01, ***p< 0.001 **(C)** Similar to **B**, but showing spontaneous measurements taken prior to exercise. Additionally, significant differences from the final measurement (day 25), are indicated as ##p < 0.01, ###p < 0.001. **(D-F)** Measurements of behavioral in the postoperative recovery period, with mice divided according only to pre-OHDA treatment (Exercise or Sedentary) including net ipsiversive rotation **(D)** absolute rotational speed **(E)**, and translational movement **(F)** indicating difference between groups by Welch t-test using *p< 0.05 **p< 0.01 ***p< 0.001.

To assess behavior without amphetamine exacerbation, we also performed recordings prior to each exercise session (Fig 2C). In the pre-lesion period (through day −1), there was no difference between mice that were exercised and those that were sedentary, nor any change over time or interaction effect (Mixed ANOVA). In the peri-lesion period (day −1 through 25), there remained no difference between treatment groups or interaction effect, but there was a significant change over time (p=3.8×10^−19^, Mixed ANOVA). We compared all post-lesion measurements to the pre-lesion baseline as well as the final measurement, to assess both hemiparkinsonian behavior and recovery. Ipsiversive rotations were significantly higher on each day post lesion, compared to day −1 (p<.05), and significantly higher on days 1, 2, and 22 compared to day 25 (p<0.01, paired t-tests, with Bonferroni correction).

Additionally, we performed recordings of the mice during the post-operative recovery from anesthesia, where mice often immediately exhibit obvious ipsiversive turning tendency. These recordings were analyzed including all mice (even those that did not reach the endpoint) and divided only according to pre-OHDA treatment, increasing statistical power. Mice that had been exercised had significantly higher net ipsiversive rotations (p=0.015, Welch t-test, Fig 2D). However, they also had a higher absolute rotational speed, including rotation in both directions (p=7.1×10^−4^, Fig 2E), as well as a higher translational movement speed (p=5.4×10^−3^, Fig 2F).

### Exploratory Behavioral Analysis

As our video recordings of open field movement allowed for metrics beyond net rotational velocity, we did an exploratory analysis to see if there were any trends toward group-wise differences. We identified behavioral features encompassing translational movement, place preference, posture, limb movement, and independent measures of ipsiversive and contraversive rotation. Considering each measure separately, we analyzed the sources of variance (Mixed ANOVA). Without controlling for type 1 error, among variables with a significant effect of measurement day relative to the lesion (uncorrected p<0.05), we assessed the effects of treatment group and the interaction of group and day (Fig 3A). Movement speed of the mouse across the field and mean head angle showed an effect (uncorrected p<0.05). While there was no clear trend in the latter, there was an apparent trend toward higher movement speed in the Exercise-Before group (Fig 3B), which aligns with the assessment of post-operative recovery (Fig 2F). However, this trend was no longer appreciable in later stages.

**Figure 3.**
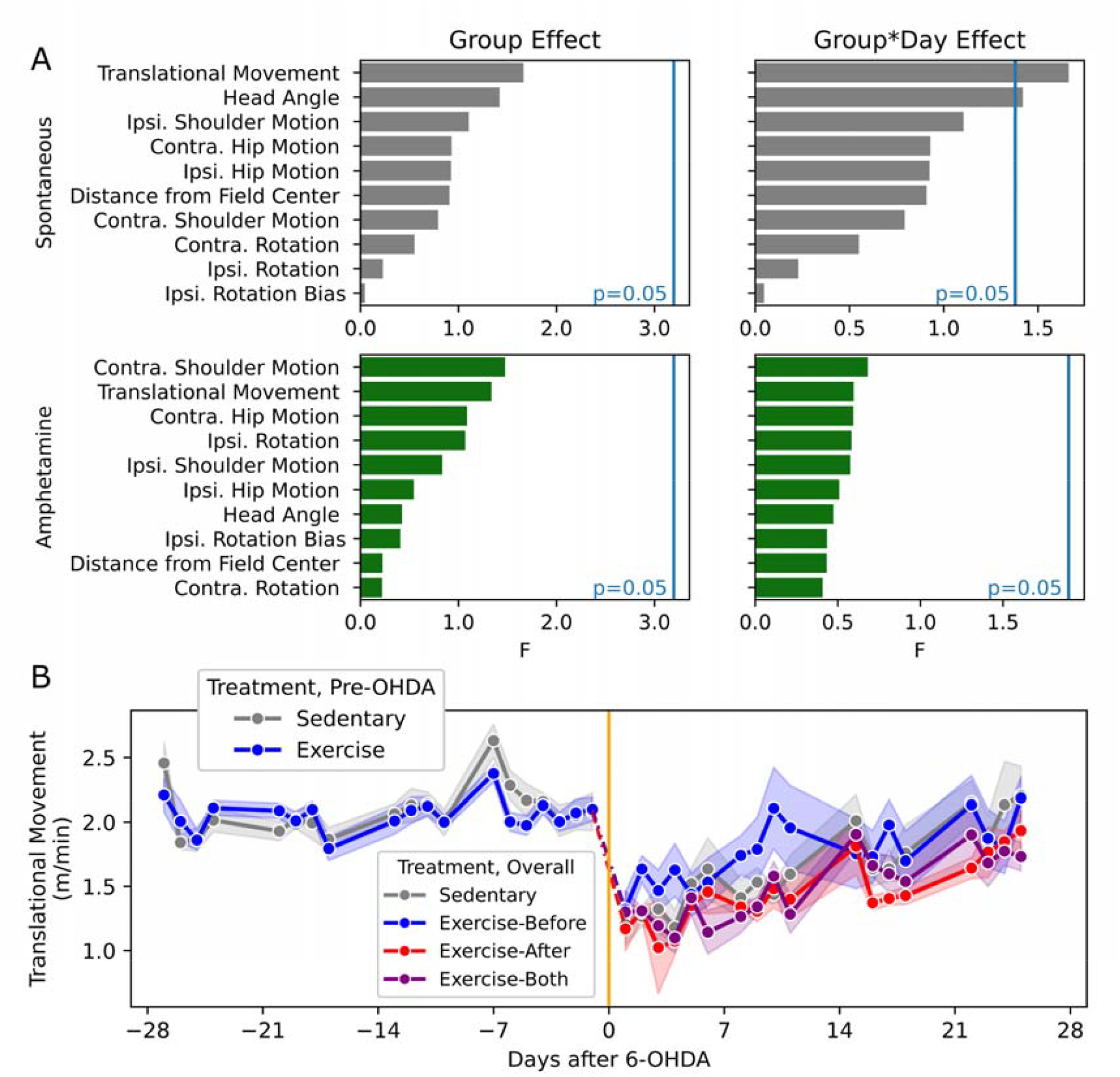
Exploratory Behavioral Measurements. **(A)** Charts showing F value for the effect of treatment group and group*time interaction in mixed ANOVA for the listed behavioral outcome variables. Vertical blue line indicates the F values where uncorrected p=0.05. **(B)** Plot of spontaneous centroid movement, showing groupwise mean (dots) and SEM (bands).

### Exercise and the Nigral Dopaminergic Cell Population

At the end of the experiment, nigrostriatal integrity was assessed by tyrosine hydroxylase (TH) immunohistochemistry in coronal sections of the midbrain and striatum (Fig 4A). Our primary measure of neuroprotection was the dopaminergic cell deficit in the substantia nigra compacta (SNc), which was measured as a ratio of the 6-OHDA injected side cell counts to the contralateral side (Fig 4B). With one exception, the lesioned side cell population was lower (Fig 4C), at a mean of 29% of the contralateral. However, there was no difference between groups (p=0.48, One-Way ANOVA, Fig 4D).

**Figure 4.**
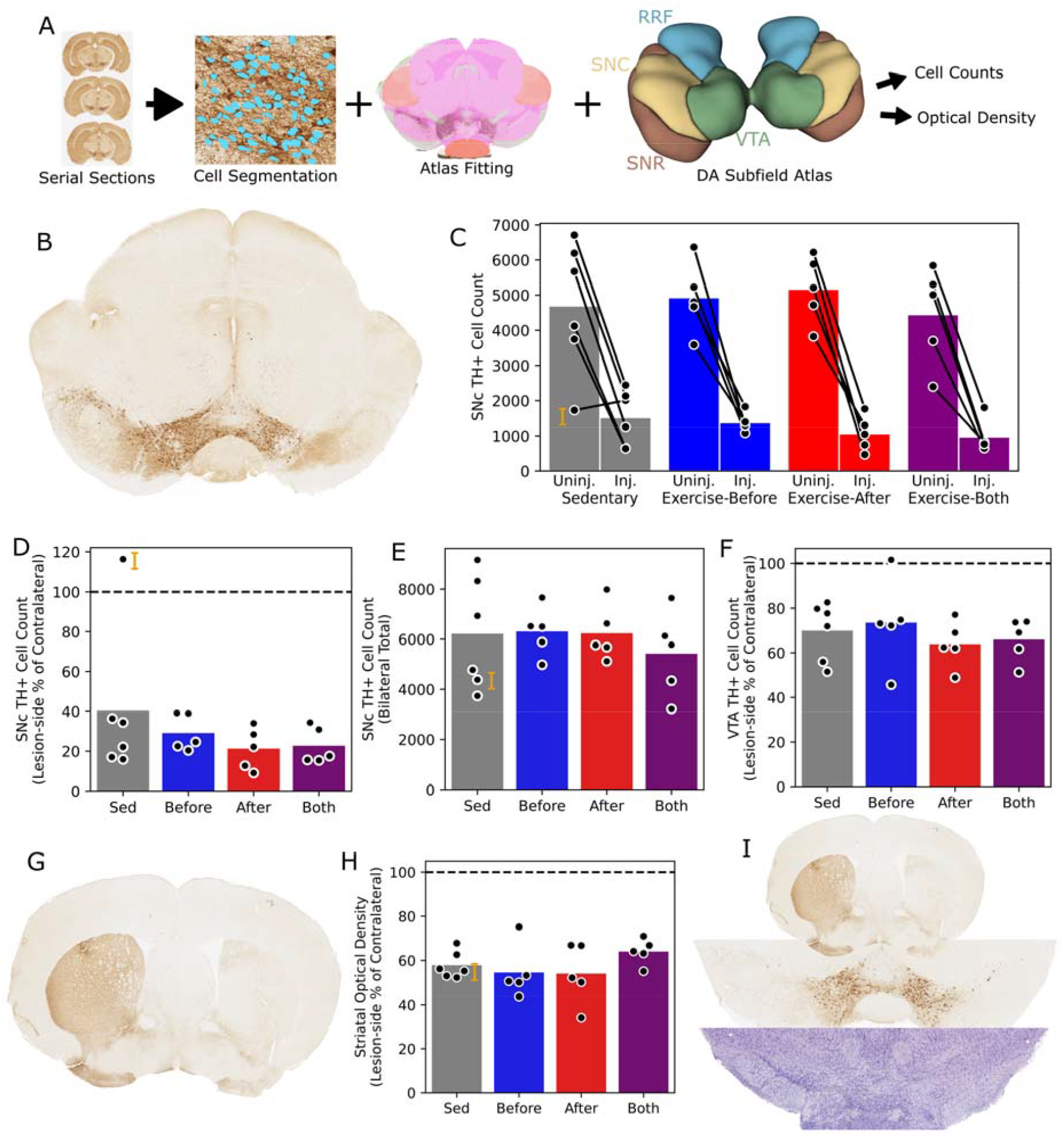
Nigrostriatal Integrity. **(A)** Chart showing histologic analysis pipeline based on TH IHC. **(B)** Representative example of lesion-side (shown on the right), TH+ cell depletion in the SNc. **(C)** Plot of SNc TH+ cell counts, divided hemisphere, showing each subject (dots and lines) and group means (bars). An outlier is indicated by the orange letter J, and shown in more detail below. **(D)** Plot of SNc TH+ cell ratios, expressed as a percentage of lesion-side cells relative to the contralateral side. **(E)** Plot of SNc TH+ cell counts, showing the bilateral sum. **(F)** Plot of VTA TH+ cell ratios. **(G)** Representative example of ipsilateral TH+ projection depletion in the striatum. **(H)** Plot of striatal TH IHC optical density, expressed as a percentage of the lesion-side mean relative to the contralateral side. **(I)** Sections from the outlier indicated elsewhere by the letter J, showing bilateral TH+ cell deficiency (middle), despite solely ipsilateral TH staining deficit in the striatum (top) and solely ipsilateral cell deficit on Nissl stain (bottom)

Given that the tyrosine hydroxylase immunopositive (TH+) cell population is plastic even in healthy animals (Aumann, 2016), it is plausible that a bilateral intervention like exercise could have contralateral or poorly lateralized neurotrophic effects. To rule this out, we also compared the total TH+ cell count between mice, again finding no difference between groups (p=0.80, Fig 4E). We also assessed the TH+ cell population in other midbrain regions, including the substantia nigra reticulata (SNr), ventral tegmental area (VTA), and retrorubral field, finding no significant group-wise differences. As expected in this model, the VTA was relatively spared by the lesion with a mean ipsilateral cell ratio of 68% (Fig 4F). Additionally, the striatal sections (Fig 4G) were assessed according to the mean optical density within the bounds of the striatum on the lesioned side, relative to the contralateral. Again, there was no difference between groups (p=0.38, Fig 4H).

In a single subject, the TH+ cell count was higher on the 6-OHDA injected side. Rather than a failed lesion, this actually reflected a bilateral TH+ cell deficit, though not a true nigrostriatal deficiency. Unlike the injected side, striatal staining did not show TH depletion of the non-injected side, and Nissl staining support did not reflect bilateral cell loss (Fig 4I).

### Dopamine Dependence of Exercise

As exercise did not abate TH+ cell loss nor the associated chronic motor deficits in 6-OHDA lesioned mice, it is notable that these mice were able to perform intensive running without obvious impairment. With this in mind, we probed the dopamine dependence of exercise, starting with a re-examination of our earlier results.

First, within the group that exercised both before and after 6-OHDA lesioning, we examined within-subject running performance across the disease process. Compared to the pre-lesion mean, mice ran significantly less on pre-lesion day −28 (the first day of exercise) as well as post-lesion days 1-8 (with day 7 being a rest day) (p<0.05, Mixed LM, Fig 5A). There was no difference thereafter, including in the later periods where dopaminergic cell loss should be well established.

**Figure 5.**
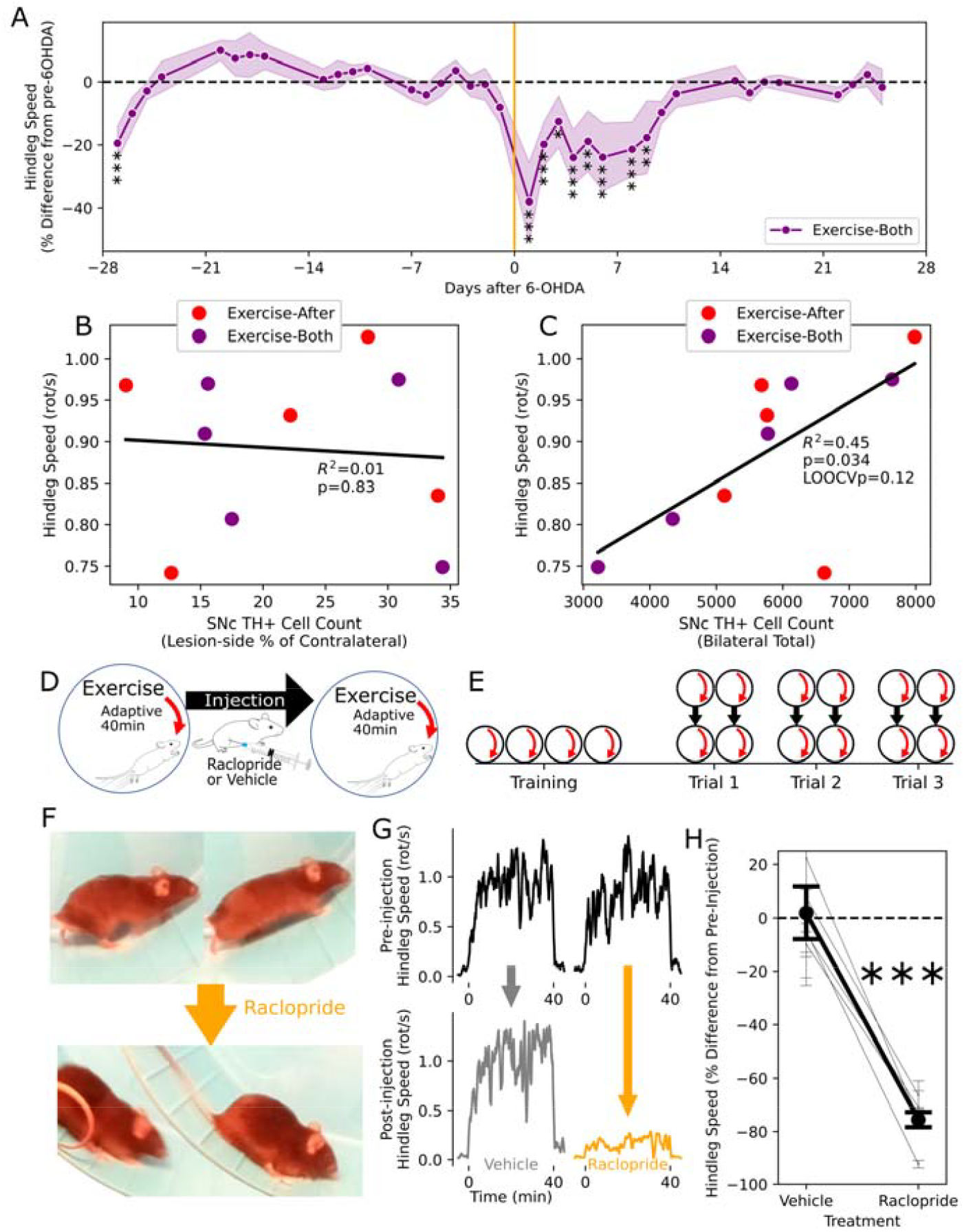
Dopamine Dependence of Exercise. **(A)** Running performance among animals exercised both before and after the 6-OHDA lesion, expressed as the mean (dots) and SEM (bands) of the percent difference from the subject-wise pre-OHDA means. Mixed LM, *p < 0.05; **p < 0.01; ***p < 0.001. **(B-C)** The relationship between exercise performance in the final week of the intervention and the dopaminergic cell population, according to the lesion side cell deficit in **B** and the total cell count in **C. (D-E)** Schematics of trial **(D)** and experimental **(E)** design for the test of D2 receptor blockade on exercise in healthy mice. **(F)** Representative images showing exercise performance before and immediately after injection with raclopride. **(G)** Representative traces showing running speed before and after injection with saline or raclopride. **(H)** Aggregate running performance after injection with vehicle and raclopride, expressed as the within-trial percentage difference, showing the overall means (heavy line) and subject means (fine lines) with accompanying SEM (error bars). Paired t-test of subject means, ***p < 0.001.

Next, within all mice that exercised after 6-OHDA lesioning, we tested the relationship between the surviving dopaminergic cell population and running performance, with the latter averaged across the final four days of exercise. Our primary measure of hemilateral dopamine depletion, the ratio of lesion to control side TH+ cells, was poorly correlated with hindleg motion during adaptive exercise (R^2^=0.01, p=0.83, linear regression, Fig 5B). However, the total count of SNc dopaminergic cells was much more predictive of running performance (R^2^=0.45, p=0.034, Fig 5C), though the relationship was not robust to leave-one-out cross-validation (p=0.12).

Finally, to determine whether running exercise was more generally dopamine dependent, we tested the effect of systemic perturbation of dopaminergic signaling with raclopride, a D_2_ receptor antagonist, in a new group of healthy mice (n=5). After an initial training period of daily exercise, mice underwent trials that entailed a baseline exercise session, followed by injection with raclopride (or a saline vehicle control), and a subsequent test exercise session (Fig 5D,E). Raclopride treatment markedly suppressed running, with mice primarily sliding down the wheel (Fig 5F), along with occasional jumping, without ever appearing sedated or immobilized during wheel motion. The post-injection performance was measured relative to the pre-injection session, being reduced by 76% after raclopride injection, compared to a 2% increase after vehicle, which was a significant difference (p=3.0×10^−4^, paired t-test of subject-wise trial means, Fig 5H).

### Exercise Physiology in the Dopamine Depleted SNr

Given that exercise evoked vigorous symmetric motor function in mice with chronic lateralizing motor deficits, without neuroprotection or lasting motor improvement, we hypothesized that it might be transiently modulating basal ganglia function towards a more functional state. To test this, we measured SNr physiology during exercise in mice with established 6-OHDA lesions. The SNr was targeted due to being the primary output nucleus of the mouse basal ganglia with well described pathophysiology after 6-OHDA lesion (Whalen et al., 2020).

Each subject (n=4) was prepared with implantation of a multi-tetrode microdrive, followed by initial exercise training without a tether (3-4 sessions), before trials commenced. Starting at the dorsal SNr, we advanced the microdrive to record new units for each repeated trial, until reaching the ventral limit of the SNr (Fig 6A). Trials were separated by at least six hours. The first set of experiments involved either adaptive exercise (40min) or a sedentary (locked wheel) control, both of which were preceded by a “Baseline” period and followed by a “Post Trial” period, which were equal in length and had the wheel locked. Despite the addition of an implant, headstage, and tether, mice consistently ran vigorously during exercise, as in prior experiments (Fig 6B,C). We isolated single-units, and limited our analysis to putative GABAergic units based on a minimum firing rate (>15Hz), with an overall mean of 35Hz in the baseline period (SD 17Hz). All analyses of neuronal firing properties were considered relative to the baseline period.

**Figure 6.**
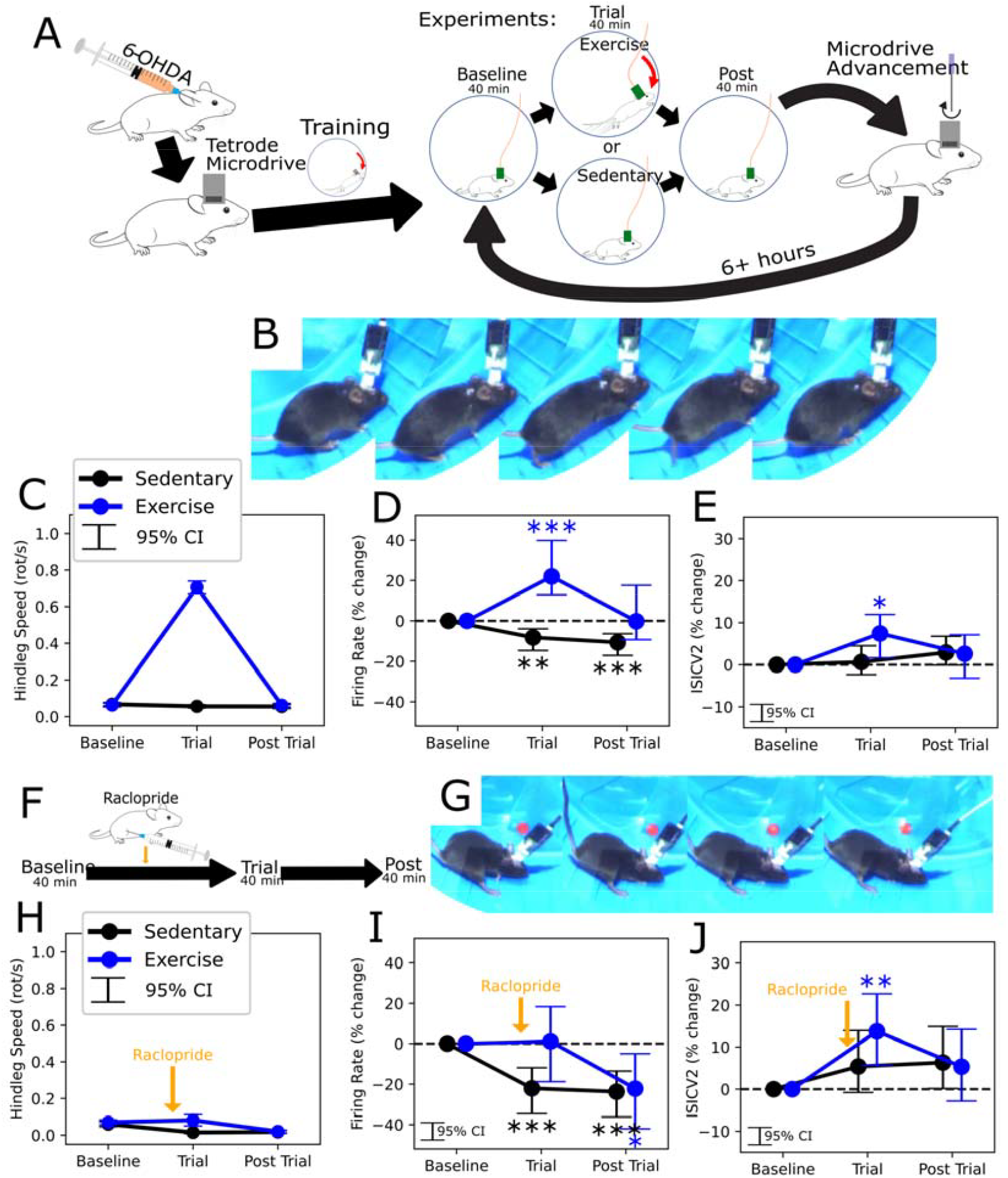
Neurophysiology of Exercise in 6-OHDA Mice. **(A)** Schematic of the experimental design, showing animal preparation, exercise (and control) trials, and electrode advancement between sets of trials. **(B)** Example of running exercise with tethered recordings, showing 1/20 consecutive frames (at 250 fps). **(C)** Aggregate running performance during exercise, measured according to hindlimb motion. **(D-E)** Changes from Baseline in firing rate **(D)** and ISICV2 during and after trials of exercise and sedentary controls. **(F)** Schematic showing addition of raclopride injection prior to the trial epoch. **(G)** Example of failure to run after raclopride injection, showing 1/20 consecutive frames. **(H-J)** Similar to **C-E**, for experiments with pre-Trial injection of raclopride as per **F**. Epochal changes were assessed via independent, linear mixed linear effect for each treatment, with stars reflecting Bonferroni corrected p values *p < 0.05; **p < 0.01; ***p < 0.001.

Exercise (n=129 units) was associated with an increase in firing rate (mean +22%, 95% CI [13,40], p=2.5×10^−4^, Fig 6D) and irregularity, measured according to a rate-insensitive coefficient of variation (CV2(Holt et al., 1996)) of the interspike interval (ISI, mean +7%, 95% CI [2,12], p=1.9×10^−2^, Fig 6E). These effects were transient, being no longer present in the Post Trial period of exercise experiments. In sedentary control experiments (n=159 units), the firing rate decreased in the Trial (mean −8%, 95% CI [−4,−15], p=1.4×10^−3^) and Post Trial epoch (mean - 11%, 95% CI [−4,−15], p=1.4×10^−3^), with no significant change in irregularity.

In order to determine whether the changes in SNr physiology were dependent on the remaining dopaminergic system, we repeated the same tests with IP injection of raclopride within 5 minutes of the start of the Trial epoch (Fig 6F). Given that an identical treatment suppressed running in healthy mice, if the same was true after 6-OHDA lesion, we also expected to be able to characterize SNr physiology during wheel motion without running. Indeed, in all trials, raclopride nearly entirely suppressed running, with mice primarily sliding at the end of the track (Fig 6G, H), as well as more rare instances of jumping. In addition, exercise trials were no longer associated with a significant change in the mean firing rate, and there was a significant decrease in the Post Trial period, relative to the Baseline (mean −22%, 95% CI [−5,−42], p=2.6×10^−2^). Exercise trials (n=68 units) were still associated with a transient increase in irregularity (mean 14%, 95% CI [6,23], p=2.3×10^−3^). In sedentary control experiments with raclopride (n=92 units), as without it, there was still a decrease in firing rate across the Trial (mean −22%, 95% CI [−12,−35], p=1.2×10^−4^) and Post Trial epochs (mean −24%, 95% CI [−14,−36], p=3.3×10^−5^), and still no significant change in irregularity.

## Discussion

Adaptive wheel exercise did not protect nigral dopaminergic cells nor alleviate chronic motor deficits in the 6-OHDA mouse model of PD. Interestingly, however, the established unabated motor lesion also did not inhibit running performance. Wheel running, in the partially dopamine depleted mice, remained sensitive to perturbation of the dopaminergic system and was associated with characteristic, transient changes in the physiology of the SNr.

### Neuroprotection with Exercise

In this study, exercise along any time course unequivocally failed to protect against nigrostriatal degeneration. Compared to sedentary mice, there was no suggestion of improved TH+ cell preservation nor a reduction in hemiparkinsonian behavioral asymmetry in any of the exercise groups. Even with an expansive analysis of open field behavior, the only detectable group difference was greater movement speed among pre-exercised mice immediately after 6-OHDA, which did not persist to the endpoint.

While unexpected, this should not be entirely surprising. As much as exercise-based neuroprotection has been widely reported in rodent models of PD, it is far from a universal finding. In fact, some studies do not show an effect on the TH+ cell population, even where therapeutic effects on motor behavior are evident (Ferreira et al., 2020; Palasz et al., 2019). Across a heterogeneous range of interventions with inconsistent outcomes, there are only limited trends suggesting which features of exercise and disease are associated with neuroprotection.

In our study, we can rule out that neuroprotection was limited by abject failure to exercise or confounded by overt aversive stimuli, though we did not assess exercise outside of measuring movement (such as by metabolic or stress indicators). However, it is worth considering that reduced running in the days immediately after the lesion may have limited neuroprotection. This early period has been directly shown to be critical for neuroprotection in a study of forced limb use in 6-OHDA rats (Tillerson et al., 2001), and more delayed running interventions tend to not to be protective (Ferreira et al., 2020). In 6-OHDA mice, dopaminergic cell loss is established, though incomplete, by nine days (Stott and Barker, 2014). As it pertains to both motor and non-motor function, including the combination of both that effects exercise, mice are particularly impaired in that early period. It may be that the severity of 6-OHDA intoxication in this species either outpaces the protective capacity of exercise or precludes effective exercise in the critical period, and that less rapid or severe models would allow for neuroprotection.

However, there are two reports of effective exercise-based protection of the SNc dopaminergic cells in 6-OHDA mice, both using the C57Bl6 strain. One applied non-aversive exercise at a slightly higher intensity than our intervention (according to track speed) for six weeks only prior to 6-OHDA injection (Aguiar et al., 2016). The other study used treadmill exercise at a higher intensity, only initiating it 7 days after the lesion (Wang et al., 2022). Both groups injected 6-OHDA in the striatum, with the former using a higher dose than our protocol, and the latter not specifying but using a higher injection volume. Both studies reported less severe lesions among their sedentary controls than our experience. Among the few other exercise studies in this animal model (Ebrahimi-Ghiri et al., 2021; Goes et al., 2014), none directly measure and report on the dopaminergic cell population, though at least one asserts a lack of dopaminergic neuroprotection (Aguiar et al., 2013).

In that context, our findings only support the established inconsistency of exercise-based neuroprotection in rodent models of PD, and certainly do not undermine the possibility that exercise has neuroprotective properties in PD in humans. Despite the advantages of adaptive wheel exercise and the 6-OHDA mouse model of PD, their combination was not suited for studying neuroprotection.

### Transient Functional Restoration with Exercise

Though it did not rescue the nigrostriatal tract, adaptive wheel exercise was able to consistently evoke running in 6-OHDA mice. As wheel running entailed vigorous, symmetric, coordinated, and partially self-paced movement, it reflects a surprising level of motor function in animals with severe unilateral lesions to the motor system and associated motor deficits. Remarkably, outside of the early post-OHDA period, running performance was not impaired at all by the dopaminergic lesion. This prompted us to explore the underlying mechanisms.

We first assessed the role of the remaining dopaminergic system, and whether wheel running was fully independent of that system either in health or after adaptation to a neurotoxic lesion. It clearly was not, being dramatically suppressed by pharmacologic D_2_ antagonism in both healthy and 6-OHDA lesioned mice, as well being transiently inhibited in the early period of 6-OHDA intoxication. Compared to 6-OHDA’s permanent, unilateral, anatomically biased, partial lesion, the raclopride treatment was brief, systemic, and D_2_-selective. Wheel running, including the retained running performance after dopamine depletion, evidently depends on some combination of the remaining dopaminergic cells, long-term adaptation, and balance between receptor-type subsystems.

Our second question was the underlying neurophysiology of exercise in the dopamine depleted basal ganglia. Measured according to single-unit activity at the outflow in the SNr, we found that exercise had a clear signature of more frequent and irregular firing. The irregularity induced by the exercise program survived D_2_ blockade, but the rate increase and running itself did not.

While the causal relationship between physiology and running is not tested here, effective running exercise was associated with a distinct, transient state of basal ganglia activity.

Though SNr physiology was measured only as a high-level aggregate in relation to a single behavioral task (running), the correlation between movement and hyperactivity is interesting, as the rate model (Albin et al., 1989; DeLong, 1990) of the basal ganglia suggests that increased activity in the output nuclei should be anti-kinetic. However, there is evidence for physiological(Wichmann, 2019; Willard et al., 2019) and anatomical(Wichmann et al., 1999) complexity in this regard, and that extends to the effects of antiparkinsonian therapies.

Suppression or ablation of the SNr (Wichmann et al., 2001; Zapletal, 1965) or homologous globus pallidus internus (GPI) (Baron et al., 2002; Laitinen et al., 1992) improves motor function in PD patients and animal models. However, subthalamic nucleus (STN) DBS, which has comparable benefits, has been shown to increase aggregate SNr activity in both humans(Galati et al., 2006) and mice (Schor et al., 2022). In monkeys, STN DBS has been shown to be similarly excitatory in the GPi (Hashimoto et al., 2003), which may be more relevant to human motor symptoms. Evidently, there are many physiological avenues to suppress the motor symptoms of PD, though the established approaches likely act more by interrupting pathophysiology than restoring functional neurophysiology. Exercise represents a distinct, naturalistic, and functional route that is worthy of further study, as it may be replicable with neuromodulation as a therapy.

### Retained Motor Function in Parkinson’s Disease

Latent, retained motor capacity is well described in PD, notably in the form of paradoxical kinesia, where patients experience a transient, contextually driven improvement in motor function. While often described in the context of stress or danger, this phenomenon can be instigated in the normal course of physical activity (Duysens and Nonnekes, 2021), including cycling (Snijders and Bloem, 2010), imagining cycling (Kikuchi et al., 2014), and catching a ball (Majsak et al., 2008). As a more explicit clinical intervention, “forced” exercise in PD patients using tandem bicycling has been shown to elicit acute improvement in motor function alongside changes in functional neuroimaging (Alberts et al., 2011; Segura et al., 2020).

Previous studies in rats have investigated paradoxical kinesia as invoked by forced swimming, where it survived dopamine blockade (Keefe et al., 1989), and by both auditory stimulus (Tonelli et al., 2018) and electrical stimulation of the inferior colliculus (Melo-Thomas and Thomas, 2015). Running exercise could provide a more translationally relevant model to study retained motor function in PD models and patients.

Paradoxical kinesia may result from temporary improvements in basal ganglia function (possibly due to evoked dopamine release) or from the takeover of motor control by alternative motor pathways, likely driven by sensory input (Melo-Thomas and Schwarting, 2023). The latter possibility may explain why the SNr hyperactivity that we observed was not anti-kinetic, and it suggests that future studies in PD models should expand upon this work to assess exercise physiology outside of the SNr.

### Lasting Effects of Exercise

Though we assert that running exercise evokes a functional, non-hemiparkinsonian state worthy of study, our results provide no evidence that performing exercise had any lasting or out-of-domain benefits, and the associated excitatory physiological changes were distinctly transient. This may be a measurement limitation imposed by our model and the simplistic metrics of spontaneous and amphetamine induced activity, which do not necessarily get at the improvements in gait (Intzandt et al., 2018; Mehrholz et al., 2015; Shen et al., 2016) and cognition (David et al., 2015; Johansson et al., 2022) that have been ascribed to exercise.

There is an element of longitudinal adaptation, in that mice gradually increased their running performance (over 2-4 days, in health), which may merit further study.

In PD patients, clinical trials of exercise have had lasting therapeutic effects. This may reflect neuroprotection, with exercise impeding nigrostriatal degeneration, which would accord with studies that show slowing but not reversal of motor symptom progression (Kolk et al., 2019; Schenkman et al., 2018; Tsukita et al., 2022). This also aligns opposite our own results, where there was a lack of neuroprotection and in turn no chronic behavioral effects.

However, exercise has also been shown to *improve* motor function (Prodoehl et al., 2015; Ridgel et al., 2009; Tollár et al., 2018), which is more difficult to ascribe to a slowing of degeneration. Though outright nigrostriatal restoration is not inconceivable, as an alternative, recent imaging studies of participants in exercise interventions have shown neuroplasticity only outside of the midbrain (Johansson et al., 2022). Exercise-based plasticity has also been demonstrated in other structures in rodents, notably the pedunculopontine nucleus (Li and Spitzer, 2020), as well as aspects of dopamine handling that are not reflected in the dopaminergic cell count (Sconce et al., 2015). While possibly involved in retained motor function after dopamine depletion, these chronic adaptations canno*t* fully explain the acute effects shown in human studies of exercise (Segura et al., 2020), cases of paradoxical kinesia, or our own results.

## Conclusion

Overall, we found that exercise was not neuroprotective in our model system, which aligns with the heterogeneous outcomes of similar studies. However, in the absence of neuroprotection, the capacity to evoke unencumbered motor function in conjunction with a characteristic neurophysiological state suggests an alternate direction for exercise studies in PD. The physiological state induced by exercise may serve as a target for improved neuromodulatory therapies, and the underlying mechanisms may offer routes to reach it. Future work should explore the dopamine dependence and SNr physiology of exercise in greater detail, in addition to the role of other systems and the potential role of chronic neuroplastic changes outside of the SNc.

## Acknowledgements

This work was supported by funding from the Department of Defense Congressionally Directed Medical Research Programs (Award no. W81XWH-19-1-0776).

## Methods

### Animals

Male C57Bl/6J strain mice (Jackson Labs) were used for all experiments. This work was performed in accordance with animal-use protocols approved by the Emory University Institutional Animal Care and Use Committee.

### 6-OHDA Injection Surgery

Hemiparkinsonism was induced by unilateral intrastriatal injection of 6-OHDA hydrobromide (Sigma-Aldrich #162957) at 16 weeks of age. To prepare for surgery, mice were anesthetized with isoflurane, given IP meloxicam 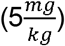 for analgesia, shaved, and fixed in a stereotactic frame (Kopf Instruments). After leveling the skull, a craniotomy was created at 1.8mm lateral and 0.4mm anterior to bregma. A glass needle in a stereotactically held micro injector (Drummond Nanoject 3) was placed at this target, advanced to a depth 3.25mm ventral from the dural surface, and used to inject 1µL of 6-OHDA hydrobromide solution (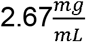 free-base, in 0.9% saline + 0.2% ascorbic acid). After injection, the needle was withdrawn, followed by wound closure and recovery. For two weeks following surgery, animals were weighed daily and provided supplementary high-calorie food and intraperitoneal fluids as needed.

### Exercise

We used adaptive wheel exercise, as previously described(Skelton et al., n.d.), for all exercise sessions. Each session included 40 minutes of exercise and took place during the light cycle. In brief, the adaptive exercise program uses color segmentation to identify mouse position and positional change as an indicator of exercise performance, with a control strategy designed to accelerate mice who are advancing along the wheel, but to rapidly decelerate if they fall off pace to permit recovery. Sedentary control sessions entailed placement in a locked wheel for the same length of time. All sessions were video recorded, and DeepLabCut (Mathis et al., 2018) was used with a custom model to identify body part coordinates. The hip joint and forward facing hindfoot coordinates were used to calculate hindleg angle. The framewise change in the hindleg angle, measured at the 90Hz resolution of the video, was used as the measure of running, with the total change averaged over the session length.

### Behavioral Assessment

We measured motor behavior using open field testing. This was done in a dark circular (225mm diameter) chamber using top-down, infrared videography. Recordings were performed prior to every exercise session (15min duration), approximately weekly (on non-exercise days) with prior injection of amphetamine (2700min duration), and in the recovery period after 6-OHDA surgery (115min duration). These recordings were processed using DeepLabCut (Mathis et al., 2018) with a custom model to identify body part coordinates. A line from the neck to hip joint was used to identify the angle of the mouse body, with angular change considered as rotation. The body center coordinates were used to determine place preference (relative to the center of the field) and movement speed.

### Drugs

To aggravate hemiparkinsonism for behavioral assessment, mice were given methamphetamine by intraperitoneal (IP) injection (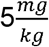, dissolved in sterile saline), immediately prior to assessment. For testing systemic perturbation of dopamine signaling with D_2_ receptor antagonism, mice were given raclopride by IP injection (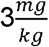, dissolved in sterile saline), with control experiments using an equivalent volume of saline vehicle, within 5 minutes prior to initiating exercise (or fixed-wheel control) trials.

### Electrophysiology Preparation and Setup

At least two weeks after 6-OHDA lesion, mice were implanted with custom fabricated, 32-channel tetrode microdrives based on a modified version of the TetrODrive (Brosch et al., 2021). The fiber optic was not used, except to provide rigidity. These were used to record electrophysiology in the running wheel (modified with electromagnetic shielding) alongside high-speed (250Hz) video capture (Teledyne FLIR Model #BFS-U3-04S2C-C). Electrophysiologic signals were amplified using a digital headstage (Intan Technologies, Part #C3324), which fed data into a USB interface board (Intan Technologies, Part #C3100) alongside shutter signals (for video synchronization), and readouts from the running wheel encoder. All components of recording and experimental control were run using custom workflows implemented with the Bonsai framework(Lopes et al., 2015).

### Electrophysiology Experiments

After preparation with the tetrode microdrive implant, mice were acclimated to exercise without a tether, before beginning electrophysiology experiments. After attachment to the tethered headstage and placement in the wheel, each recording session consisted of a Baseline, Trial, and Post Trial period, each lasting 40 minutes. The Trial period consisted of adaptive exercise, as used elsewhere, or a sedentary control condition with a locked wheel. In the trials with raclopride, the mouse was removed from the wheel within 5 minutes before the onset of the Trial period for IP injection, and promptly returned to the wheel. The period outside of the wheel was identified on video and excluded from analyses. We waited at least 6 hours between each trial, and advanced the microdrive (125µm) to record from new units before repeating any set of experimental conditions, recording from 8-9 positions for each animal. With each recording depth, experimental conditions alternated between exercise and sedentary, with or without raclopride, with the order of exercise and sedentary experiments reversed for each advancement.

### Electrophysiology Analysis

After preprocessing using a bandpass filter (600 to 6000Hz) and common median referencing within tetrodes, units were identified using Kilosort2 (Pachitariu et al., 2016) within SpikeInterface (Buccino et al., 2020) and post-processed with the automatic merging algorithm from Lussac (Llobet et al., 2022). Isolated single units were identified based on the prediction provided by Kilosort in addition to having an ISI violation ratio less than 0.1 and signal-to-noise ratio greater than 4. Dopaminergic units were excluded by a minimum firing rate threshold (>15Hz), and the remainder were treated as GABAergic. Unit firing properties were assessed according to the firing rate and the irregularity, which was measured using the CV2 rate invariant coefficient of variation (Holt et al., 1996) of the ISI. The exercise videos were analyzed as described for previous experiments, with the frame level data synchronized to the electrophysiology using the shutter signal.

### Tissue Preparation

On the study endpoint day, mice were sacrificed by IP injection of pentobarbital/phenytoin solution (0.5mL, Virbac Euthasol, 1:39 dilution of stock solution in saline) and transcardial perfusion with phosphate-buffered saline (PBS) and 4% paraformaldehyde (PFA). The brain was dissected from the skull, and after at least 24 hours in PFA was moved to 30% sucrose solution until sinking, for cryoprotection. Brains were then sectioned coronally on a cryostat at 40µm thickness, collected in 8 series, and retained as floating sections. For tyrosine hydroxylase immunohistochemistry, one out of four sections was used. To prepare for staining, sections were permeabilized (3% H_2_O_2_ + 0.1% Triton X-100, 30 minutes), rinsed (PBS, 10min x 3) and blocked (4% normal donkey serum, 30 minutes). They were then incubated in primary antibody (Rabbit anti-TH, 1:1000 dilution, Pel-Freeze P40101, 4° C, overnight), rinsed, and incubated in a biotinalyted donkey anti-rabbit secondary antibody for an hour at room temperature, rinsed, and incubated with an avidin-bound detection enzyme (Vectastain ABC-HRP Kit #PK-4001,4° C, overnight). Sections were then rinsed, and staining was completed with addition of the chromogen (3-39-diaminobenzidine tetrachloride) for 12 minutes in the presence of H_2_O_2_. After a final rinse, the sections were mounted on glass slides, dehydrated, and coverslipped.

### Tissue Analysis

All slides were digitally imaged using a slide scanner (Leica Aperio AT2). Each section was registered to the Allen Brain Atlas (Sunkin et al., 2013) space via automated registration with DeepSlice(Carey et al., 2023) followed by manual curation with Quicknii (Puchades et al., 2019). Cells were segmented using Cellpose (Stringer et al., 2021) with a custom model for TH immunohistochemistry. This was trained across an array of different samples from different subjects until it was found to consistently produce accurate segmentation in new samples. Cell coordinates were identified in section space using the geometric center of the segmentation masks, and were then transformed into Allen Brain Atlas space using the registration. Cell counts within each dopaminergic subfield were determined using an anatomical atlas in Allen space, with the number of counted cells in each segment divided by the proportion of sampled segment volume to estimate the total cell count.

### Statistics

Longitudinal behavioral measurements were analyzed using mixed ANOVA, considering the effects of experimental day and treatment group, and their interaction. Post-hoc testing used pair t-tests with Bonferroni correction for multiple comparisons. Endpoint measurements of histology were compared between groups via One-Way ANOVA. For the mice exercised before and after the lesion, within-subject exercise performance was assessed relative to the pre-OHDA mean, and compared over time using a linear mixed model with fixed effects for time and random subject-level intercepts and effects of time. For all mice exercised after the lesion, the correlation of TH+ cell counts (both relative and total) was compared to exercise performance by linear regression, with significant relationships further tested with leave-one-out cross-validation. To measure the effects of raclopride on exercise performance in healthy animals, performance was measured relative to a pre-injection session, and the subject-wise means across 3 trials were compared by paired t-test. For the electrophysiology studies, unit firing properties (relative to the baseline epoch) were assessed using separate mixed models for the exercise and sedentary trials with and without raclopride. These models included fixed effects of epoch and clustered random effects of subject, trial, and unit, with statistical significance adjusted by Bonferroni correction. All statistical analyses were performed using the Pingouin (Vallat, 2018) and Statsmodels (Seabold and Perktold, 2010) packages in Python.

